# KatG catalase deficiency confers bedaquiline hyper-susceptibility to isoniazid resistant *Mycobacterium tuberculosis*

**DOI:** 10.1101/2023.10.17.562707

**Authors:** N Ofori-Anyinam, M Hamblin, ML Coldren, B Li, G Mereddy, M Shaikh, A Shah, N Ranu, S Lu, PC Blainey, S Ma, JJ Collins, JH Yang

## Abstract

Multidrug-resistant tuberculosis (MDR-TB) is a growing source of global mortality and threatens global control of tuberculosis (TB) disease. The diarylquinoline bedaquiline (BDQ) recently emerged as a highly efficacious drug against MDR-TB, defined as resistance to the first-line drugs isoniazid (INH) and rifampin. INH resistance is primarily caused by loss-of-function mutations in the catalase KatG, but mechanisms underlying BDQ’s efficacy against MDR-TB remain unknown. Here we employ a systems biology approach to investigate BDQ hyper-susceptibility in INH-resistant *Mycobacterium tuberculosis*. We found hyper-susceptibility to BDQ in INH-resistant cells is due to several physiological changes induced by KatG deficiency, including increased susceptibility to reactive oxygen species and DNA damage, remodeling of transcriptional programs, and metabolic repression of folate biosynthesis. We demonstrate BDQ hyper-susceptibility is common in INH-resistant clinical isolates. Collectively, these results highlight how altered bacterial physiology can impact drug efficacy in drug-resistant bacteria.

## INTRODUCTION

Despite significant advances in anti-tubercular drug development over the past 80 years, tuberculosis (TB) remains the leading cause of deaths worldwide due to bacterial infection^1^. *Mycobacterium tuberculosis* (Mtb) is the causative agent underlying TB disease. Standard curative treatment for TB involves at least 4 months of treatment with regimen containing isoniazid (INH) and rifampin (RIF) or rifapentine^2, 3^ along with additional drugs. However, drug-resistant and multidrug-resistant (MDR; defined as resistance to both INH and RIF) TB pose growing threats to global TB control^1^. These pressing challenges have stirred significant drug discovery efforts for treating MDR-TB and have resulted in drugs such as bedaquiline (BDQ)^4^ which exhibits exceptional activity against MDR-TB^5^. BDQ is now a cornerstone for several MDR-TB regimens, including bedaquiline-pretomanid-linezolid-moxifloxacin (BPaLM)^6^.

INH is a prodrug that is first activated by the mycobacterial catalase-peroxidase KatG (*Rv1908c*) to form a INH-NAD adduct^7^. Activated INH-NAD then tightly binds and inhibits the enoyl-acyl carrier protein reductase InhA (*Rv1484*), which is essential for the synthesis of mycolic acids that form the mycobacterial cell wall. INH resistance is most commonly associated with loss-of-function mutations in *katG* or mutations in the promoter region of *inhA* which induces InhA over-expression^8, 9^. BDQ targets the C subunit of mycobacterial ATP synthase, encoded by *atpE* (*Rv1305*), and is proposed to exert slow mycobacterial lethality by either ATP depletion^10, 11^ or by electron transport chain uncoupling^12^. Because BDQ is now a cornerstone for treating MDR-TB, it is important to understand how and why BDQ is efficacious against MDR-TB to guide future drug discovery efforts.

Here we employed a systems biology approach to investigate how KatG deficiency enables BDQ hyper-susceptibility in INH-resistant Mtb. Several groups have previously reported that BDQ antagonizes isoniazid lethality^13–15^. Here we report unexpected synergy between INH and BDQ co-treatment in Mtb which we discovered is due to BDQ-induced suppression of INH resistance evolution. We found BDQ hyper-susceptibility in INH-resistant cells is due to several physiological changes induced by KatG deficiency, including increased susceptibility to BDQ-induced reactive oxygen species (ROS) and DNA damage, synergistic repression of *inhA* and *atpE*, remodeling of mycobacterial transcriptional programs, and metabolic repression of folate biosynthesis. Importantly, we found BDQ hyper-susceptibility is common in drug-resistant and MDR strains curated by the World Health Organization and United Nations^16, 17^. Together, these findings provide mechanistic insight into how physiological changes induced by acquired resistance may enable collateral sensitivity to other drugs and guide selection of efficacious anti-tubercular regimens for treating MDR-TB.

## RESULTS

### BDQ synergizes with INH by suppressing INH resistance evolution

Performing week-long experiments, several groups have demonstrated that electron transport inhibitors, including BDQ, antagonistically interfere with isoniazid lethality^13–15^. However, while INH lethality rapidly occurs over a period of days, BDQ exerts slow bactericidal lethality over a period of weeks^11^. We therefore sought to determine if long-term co-treatment with INH and BDQ might be synergistic or antagonistic. We co-treated Mtb H37Rv cells with 0.1 μg/mL INH (2x minimal inhibitory concentration [MIC]) and a range of BDQ concentrations in 7H9 OADC medium and measured colony forming units (CFUs) over 30 days. Consistent with previous studies^13–15^, we found BDQ suppressed early-phase lethality. However, unexpectedly and in contrast, we discovered synergistic lethality between BDQ and INH in a dose-dependent manner (Fig. 1a).

**Fig. 1.**
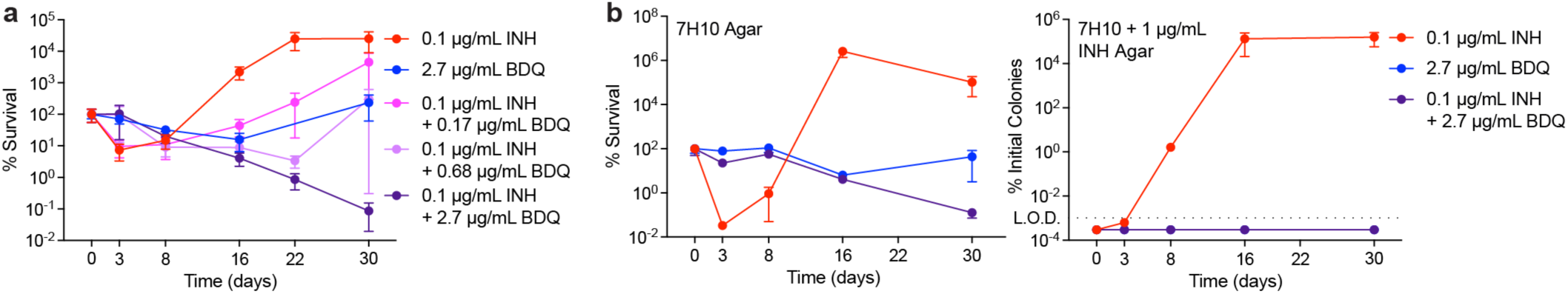
Bedaquiline suppresses emergence of isoniazid resistance. **a**, BDQ induces biphasic synergistic lethality with INH over long treatment times. Mtb H37Rv cells were treated with 0.1 μg/mL INH and varying concentrations of BDQ in 7H9 OADC medium over 30 days. **b**, BDQ suppresses emergence of INH-resistant cells. Mtb cells were treated with 0.1 μg/mL INH, 2.7 μg/mL BDQ, or their combination over 30 days and plated on 7H10 agar alone (left) or 7H10 + 1 μg/mL INH agar (right). Data shown represent % CFU normalized to CFU from Day 0 from n ≥ 3 biological replicates. Error bars represent standard errors of the mean.

INH treatment elicits rapid INH resistance *in vitro* and *in vivo*^8^. We hypothesized synergistic lethality between BDQ and INH might be explained by suppression of INH resistance formation. To test this hypothesis, we cultured H37Rv cells with 0.1 μg/mL INH alone, 2.7 μg/mL BDQ alone (which we found to be approximately bacteriostatic), or the combination of 0.1 μg/mL INH and 2.7 μg/mL BDQ (which conferred a 2- to 3-log reduction in CFU), and plated each of these treated cells on 7H10 agar plates with or without 1 μg/mL INH supplementation (Fig. 1b). Consistent with the rebound in CFU at 3 days, we found cells treated with INH alone rapidly formed CFU on INH agar plates, but cells treated with BDQ alone or INH and BDQ co-treatment did not. We reasoned the absence of colonies from co-treatment cultures on INH-containing plates indicated that BDQ suppresses INH resistance evolution and that INH-resistant cells might be hyper-susceptible to BDQ.

### Bedaquiline hyper-susceptibility is drive by a loss-of-function of KatG

We sought to test the hypothesis that INH-resistant cells were hyper-susceptible to BDQ. To select a representative laboratory-evolved INH-resistant strain that could be used for further experimental characterization, we whole-genome sequenced 6 INH-resistant colonies plated on 7H10 agar without INH supplementation after 30 days treatment with INH alone. We found that all 6 colonies harbored *katG* mutations and none possessed mutations in *inhA* or its promoter region (Supplemental Table 1). 4 of these 6 colonies possessed E553K mutations. We selected one of these to be a representative laboratory-evolved INH-resistant strain (H37Rv *katG^-^*).

To better understand how BDQ might act on INH-resistant cells, we cultured wild-type H37Rv and our laboratory-evolved H37Rv *katG^-^* in 7H9 OADC medium and quantified CFUs following treatment with 2.7 μg/mL BDQ alone (Fig. 2a). We observed a 4-log CFU reduction in *katG^-^* cells over 30 days, while CFUs remained static in wild-type cells. To confirm this phenotype was specifically due to a KatG loss-of-function, we repeated these experiments in a *katG* deletion mutant and similarly observed a 4-5-log reduction in CFU (Fig. 2b). Both these strains exhibited BDQ sensitization in dose-response growth inhibition experiments (Fig. 2c and 2d), supporting recent observations that some laboratory-evolved INH-resistant Mtb cells may possess lower BDQ MICs^18^.

**Fig. 2.**
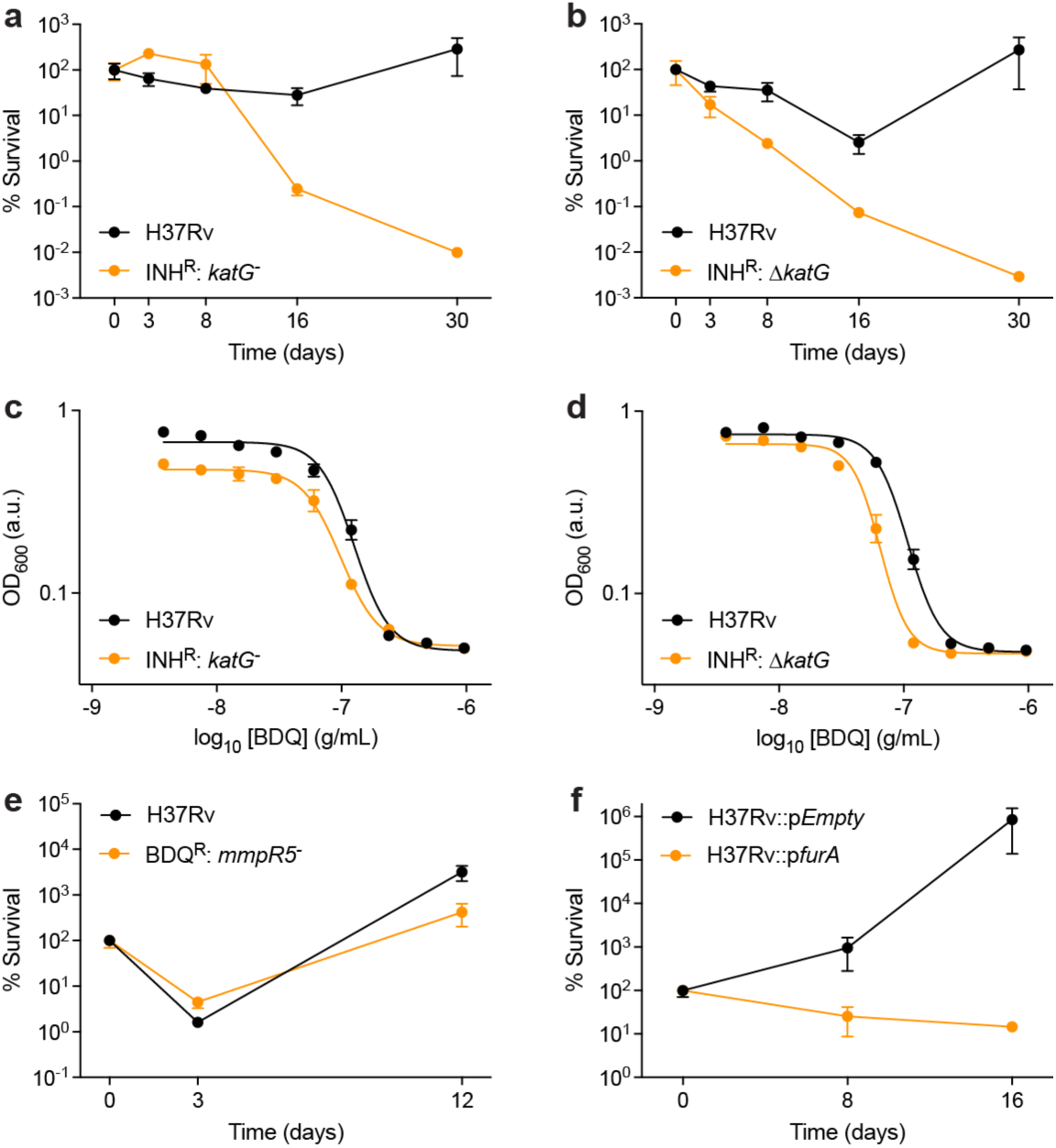
KatG deficiency underlies bedaquiline hyper-susceptibility. **a**, A laboratory-evolved INH-resistant strain harbouring a *katG* loss-of-function mutation (*katG^-^*E553K) is hyper-susceptible to 2.7 μg/mL BDQ in 30-day time-kill experiments. **b**, H37Rv Δ*katG* cells are hyper-susceptible to 2.7 μg/mL BDQ in 30-day time-kill experiments. **c**, *katG^-^* cells are modestly sensitized to BDQ in 14-day dose- response experiments. **d**, Δ*katG* cells are sensitized to BDQ in 14-day dose-response. **e**, A laboratory-evolved BDQ-resistant strain harbouring a *mmpR5* loss-of-function mutation (*mmpR5^-^* G65R) is not hyper-susceptible to 0.1 μg/mL INH in 12-day time-kill experiments. **f**, A FurA over-expression mutant is hyper-susceptible to 2.7 μg/mL BDQ in 30-day time-kill experiments. Time-kill data shown represent % CFU normalized to CFU from Day 0 from n = 3 biological replicates. Dose-response experiments represent n = 3 biological replicates. Error bars represent standard errors of the mean.

We considered the possibility synergy between INH and BDQ co-treatment might alternatively be explained by hyper-susceptibility of BDQ-resistant cells to INH. To evaluate this hypothesis, we whole-genome sequenced 5 colonies plated on 7H10 agar without INH supplementation after 30 days treatment with BDQ alone. BDQ resistance is principally governed by mutations in *mmpR5* (*Rv0678*), which represses expression of the MmpS5-MmpL5 efflux pump^19^. Sequencing revealed all 5 colonies harbored mutations in *mmpR5* and none in BDQ’s target *atpE* (Supplemental Table 1). We selected one of these to be a representative laboratory-evolved BDQ-resistant strain (H37Rv *mmpR^-^*) and treated these cells with 0.1 μg/mL INH for 12 days (Fig. 2e). We did not observe any significant differences between wild-type and *mmpR^-^* cells, indicating long-term synergy between INH and BDQ is due to hyper-susceptibility of INH-resistant cells to BDQ and not hyper-susceptibility of BDQ-resistant cells to INH.

To further validate this hypothesis, we also tested H37Rv cells transformed with a plasmid harboring an anhydrotetracycline (aTc)-inducible *furA* (*Rv1909c*) expression cassette^20–22^. FurA is a member of the ferric uptake regulator (Fur) family of bacterial transcriptional factors which represses *katG* expression in mycobacteria when up-regulated^20, 23^. We found *katG* repression induced by FurA over-expression conferred a modest 1-2-log reduction in CFU over uninduced cells over 16 days (Fig. 2f). Collectively, these results indicate KatG loss-of-function is sufficient for conferring BDQ hyper-susceptibility in INH-resistant Mtb.

### Bedaquiline potentiates reactive oxygen species formation in KatG-deficient cells

In most aerobic bacteria, the catalase-peroxidase KatG and alkyl hydroperoxide reductase AhpC mediate protection from oxidative stress by scavenging toxic H_2_O_2_^24^. Expression of these enzymes is normally regulated by the transcription factor OxyR^25^. However, the *oxyR* homologue in Mtb possesses several lesions and mutations that render it inactive and it is therefore unable to induce *ahpC* expression in response to oxidative stress^26^. Consequently, KatG is Mtb’s primary defense against oxidative stress, and KatG-deficient cells are hyper-susceptible to ROS (Fig. 3a).

**Fig. 3.**
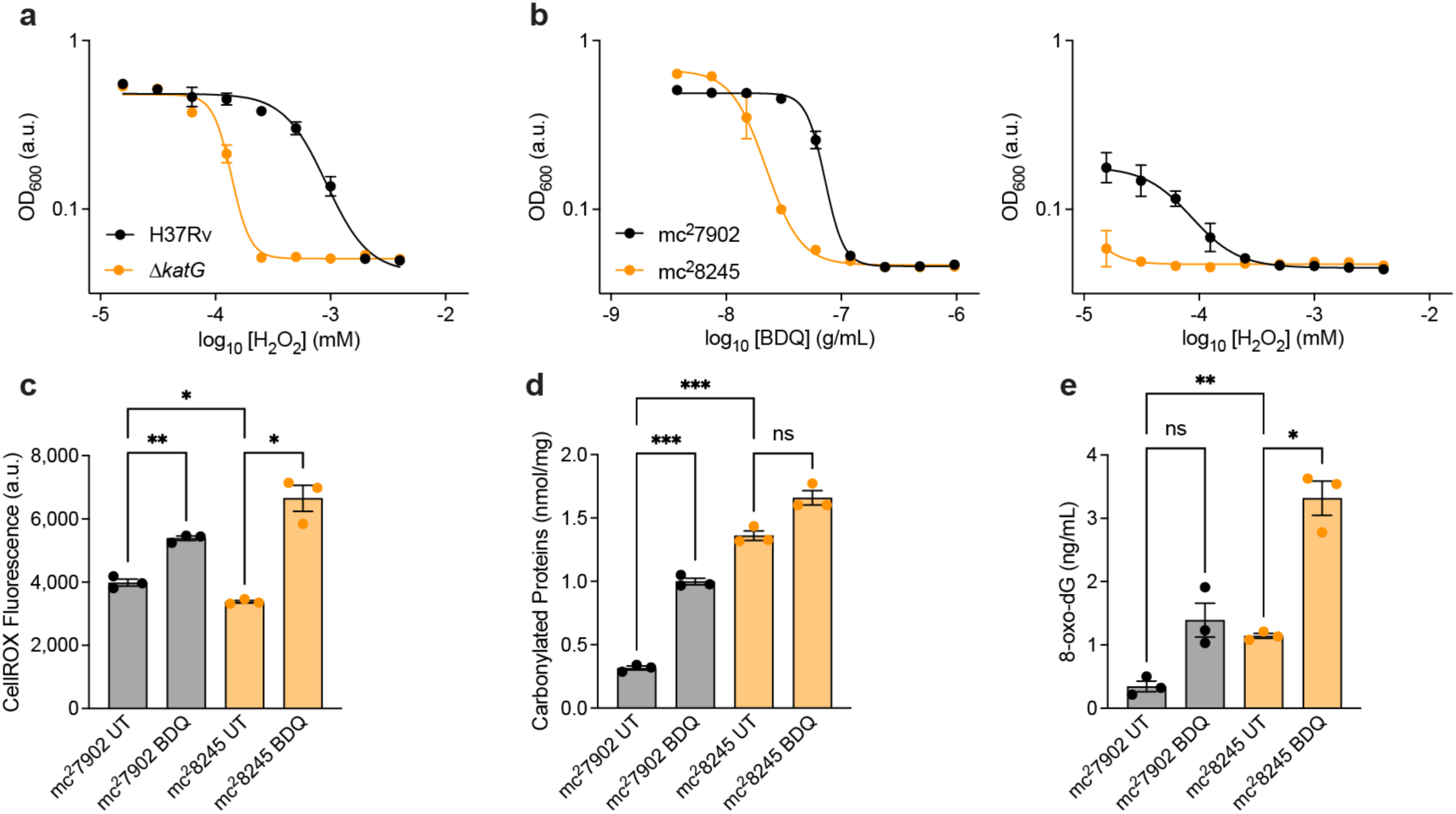
Bedaquiline potentiates reactive oxygen species formation in KatG-deficient cells. **a**, Δ*katG* cells are hyper-susceptible to H_2_O_2_ in 14-day dose-response experiments. **b**, KatG-deficient mc^2^8245 cells are hyper-susceptible to BDQ and H_2_O_2_ in 14-day dose-response experiments. **c**, BDQ induces greater increases in ROS accumulation in KatG-deficient mc^2^8245 over KatG-replete mc^2^7902 cells as reported by CellROX over 4 days treatment with 0.68 μg/mL BDQ. **d**, Basal protein carbonylation is potentiated in mc^2^8245 over mc^2^7902 cells as reported by ELISA. Protein carbonylation is potentiated in mc^2^8245 over mc^2^7902 cells after 16 hours treatment with 5.4 μg/mL BDQ. **e**, Basal guanine oxidation is potentiated in mc^2^8245 over mc^2^7902 cells as reported by ELISA. Guanine oxidation is potentiated in mc^2^8245 over mc^2^7902 cells after 16 hours treatment with 5.4 μg/mL BDQ. n = 3 biological replicates for most experiments. n = 2 biological replicates for mc^2^7902 BDQ dose-response experiments. Brown-Forsythe and Welch ANOVA statistical tests were performed on CellROX, protein carbonylation, and guanine oxidation experiments, with comparisons between BDQ-treated and untreated cells wild-type and Δ*katG* cells or between untreated wild-type and Δ*katG* as indicated with FDR correction. *: p ≤ 0.05, **: p ≤ 0.01, ***: p ≤ 0.001.

BDQ induces oxidative stress response regulons in non-tuberculosis mycobacteria^12^, but has never been directly shown to capably induce ROS formation in Mtb^27^. We hypothesized that KatG deficiency in INH-resistant Mtb would elicit BDQ-induced ROS formation. To test this hypothesis, we performed oxidative stress experiments in the INH-resistant BSL-2 auxotrophic strain mc^2^8245, which harbors a large genomic deletion spanning over *katG*, and its ancestor mc^2^7902^28^. We first validated that mc^2^8245 exhibits hyper-susceptibility to BDQ and H_2_O_2_ (Fig. 3b) over mc^2^7902. We then evaluated ROS production in these cells using the ROS-sensitive dye CellROX Green in the presence and absence of BDQ treatment. We surprisingly observed enhanced CellROX fluorescence under BDQ treatment for both strains, indicating BDQ sufficiently induces ROS accumulation in KatG-replete Mtb (Fig. 3c). Interestingly, BDQ treatment elicited a greater increase in CellROX fluorescence in mc^2^8245 cells over mc^2^7902 cells, supporting our hypothesis that KatG-deficiency potentiates BDQ-induced ROS formation. We further hypothesized elevated ROS accumulation would result in increased ROS-mediated cellular damage. To test this hypothesis, we performed enzyme-linked immunosorbent assays (ELISAs) for protein carbonylation and guanine oxidation (8-oxo-dG) in mc^2^7902 and mc^2^8245 in the presence and absence of BDQ treatment. Consistent with our CellROX measurements, we found increased baseline protein carbonylation (Fig. 3d) and accumulation of 8-oxo-dG (Fig. 3e) in mc^2^8245 cells over mc^2^7902 cells. These are consistent with expectation for KatG deficiency and reveal increased BDQ-induced oxidative damage in mc^2^8245 cells over mc^2^7902 cells. Together, these results strongly indicate BDQ induces ROS in both KatG-replete and KatG-deficient Mtb and suggest BDQ-induced ROS may contribute to BDQ hyper-susceptibility in KatG-deficient INH-resistant cells.

### Transcriptional remodeling sensitizes KatG-deficient cells to bedaquiline

To further understand how KatG deficiency enables hyper-susceptibility, we performed RNA sequencing analyses on wild-type and Δ*katG* cells with and without 2.7 μg/mL BDQ treatment for 16 hours (Fig. 4a; Supplemental Table 2). We first analyzed transcriptomic differences between BDQ-treated and untreated wild-type cells. Consistent with previous reports^11, 29^, we found BDQ induced increased expression of the general stress response regulator *devR* (*dosR*, *Rv3133c*) and several genes involved in central carbon metabolism, including genes in the ATP synthase operon (*Rv1303*-*Rv1307*), and decreased expression of ribosomal subunit genes (*Rv0703*, *Rv0719*). In addition, supporting our findings that BDQ sufficiently induces ROS (Fig. 3), we found induction of *katG* and the oxidative stress regulator *oxyS* (*Rv0117*) in wild-type cells (Fig. 4b). Interestingly, we found BDQ also decreased expression of INH’s target *inhA* (Fig. 4c).

**Fig. 4.**
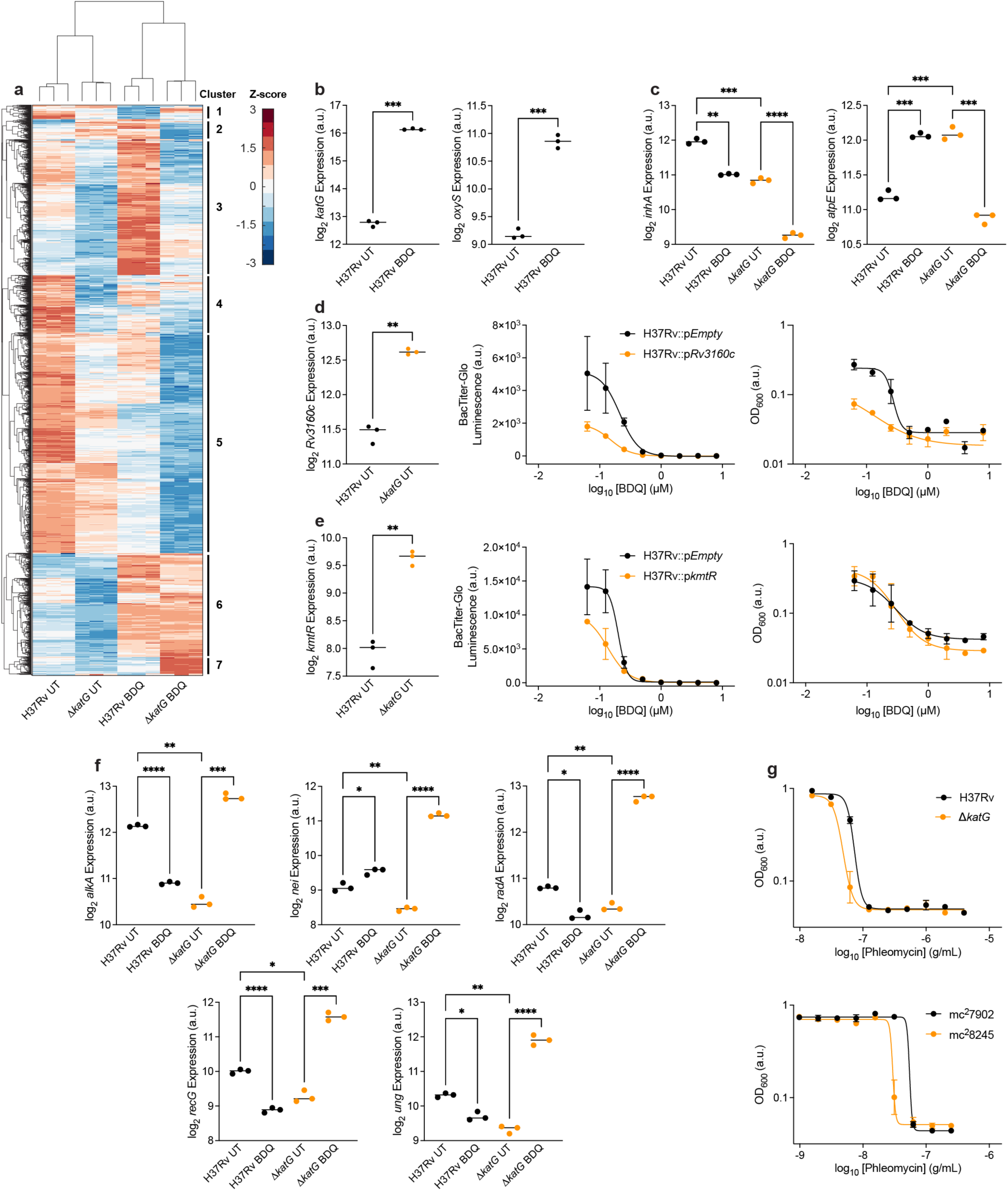
Transcriptional remodeling sensitizes KatG-deficient cells to bedaquiline. **a**, Smooth quantile normalized^73^ RNA sequencing gene expression data from BDQ-treated and untreated H37Rv and Δ*katG* cells. Clusters defined by hierarchical clustering, illustrating differences in BDQ-induced expression changes between H37Rv and Δ*katG* cells. **b**, *katG* and *oxyS* expression in untreated H37Rv and Δ*katG* cells. **c**, *inhA* and *atpE* expression in BDQ-treated and untreated H37Rv and Δ*katG* cells. **d**, *Rv3160c* expression is higher in untreated Δ*katG* cells over H37Rv cells. *Rv3160c* over-expression mutants are sensitized to BDQ in 7-day dose-response experiments by BacTiter-Glo and optical density. **e**, *kmtR* expression is higher in untreated Δ*katG* cells over H37Rv cells. *kmtR* over-expression mutants are sensitized to BDQ in 7-day dose-response experiments by BacTiter-Glo. **f**, BDQ-induced DNA repair enzyme expression is increased in Δ*katG* cells over H37Rv cells. *alkA*, *nei*, *radA*, *recG*, and *ung* are members of gene Cluster 7. **g**, Δ*katG* and mc^2^8245 cells are hyper-susceptible to the DNA damaging agent phleomycin in 14-day dose-response experiments. Smooth quantile normalized expression data given on log_2_ scales. n = 3 biological replicates for each experiment. Brown-Forsythe and Welch ANOVA statistical tests were performed on RNA expression data with comparisons between BDQ-treated and untreated cells wild-type and Δ*katG* cells or between untreated wild-type and Δ*katG* as indicated with FDR correction. *: p ≤ 0.05, **: p ≤ 0.01, ***: p ≤ 0.001, ****: p ≤ 0.0001.

We next analyzed differences between wild-type and Δ*katG* H37Rv cells. As would be expected under catalase deficiency, we observed increased expression of *devR*; the ROS detoxifying enzymes *ahpC*, *ahpD* (*Rv2429*), and *trxC* (*Rv3914*); and biosynthesis genes for the antioxidant amino acid ergothioneine (*Rv3701c-Rv3704c*) in Δ*katG* cells. Interestingly, we also observed increased expression of ATP synthase operon genes and decreased expression of several genes involved in mycolic acid biosynthesis, including *inhA*, *fabG* (*Rv1483*), *fabH* (*Rv0553c*), and *kasA* (*Rv2245*). Moreover, BDQ treatment synergized with *katG* deletion to further suppress expression of INH’s target *inhA* (Fig. 4c). We considered a similar synergy might also occur for BDQ’s target *atpE*. Interestingly, we found while BDQ induced *atpE* expression in wild-type cells, *atpE* expression was synergistically repressed in Δ*katG* cells. Together, these results suggest synergistic repression of *inhA* and/or *atpE* expression may help sensitize KatG-deficient cells to BDQ.

We previously found that Mtb remodels its transcriptional state to decrease its susceptibility to BDQ^29^. We therefore hypothesized KatG-deficient cells might also remodel their transcriptional state to alther their susceptibility to BDQ. We analyzed differences in transcription factor expression between Δ*katG* and wild-type cells and discovered 12 transcription factors significantly induced with at least 2-fold increased expression in Δ*katG* cells. These include several transcription factors previously associated with Mtb persistence or drug resistance: *devR*, *prpR* (*Rv1129c*)^30^, *blaI* (*Rv2160c*)^31^, *mprA* (*Rv0981*)^32^, *carD* (*Rv3583c*)^33^, *phoY2* (*Rv0821c*)^34^, and notably *mmpR5*, the primary resistance gene for BDQ^19^. We hypothesized induction of transcriptional programs regulated by these transcription factors might render Mtb hyper-susceptible to BDQ. To test this hypothesis, we performed BDQ dose-response experiments on empty vector control cells and H37Rv cells harboring aTc-inducible expression plasmids for the 3 transcription factors with greatest increased expression in Δ*katG* cells (*kmtR*, *prpR*, *Rv3160c*; Supplemental Table 2)^20, 21^. In support of our hypothesis, *kmtR* and *Rv3160c* over-expression strains exhibited sensitization to BDQ, whether by BacTiter-Glo or optical density (Fig. 4d and 4e). We did not observe significant differences in BDQ sensitivity in *prpR* over-expressing cells (Extended Data Fig. 1). Nonetheless, together these results suggest basal induction of hyper-sensitizing transcriptional programs may contribute to BDQ hyper-susceptibility in KatG-deficient cells.

Based on our observation that BDQ treatment and *katG* deletion synergistically suppressed *inhA* and *atpE* expression, we hypothesized other epistatic transcriptional changes might also contribute to BDQ hyper-susceptibility. To evaluate this hypothesis, we hierarchically clustered our RNA expression data and defined clusters based on patterns of BDQ-induced gene expression changes between wild-type and Δ*katG* cells (Fig. 4a; Supplemental Table 3). Most transcriptomic differences following BDQ treatment appeared shared between wild-type and Δ*katG* cells. Gene Ontology analyses revealed BDQ treatment most strongly repressed processes involved in cell wall biosynthesis, including several mycolic acid biosynthesis genes (Cluster 5: BDQ-induced down-regulation in both wild-type and Δ*katG* cells); and potentiated processes involved in cholesterol metabolism (Cluster 6: BDQ-induced up-regulation in both wild-type and Δ*katG* cells) (Supplemental Table 3).

We reasoned the most informative clusters would contain expression changes that differed between wild-type and Δ*katG* cells. Although Gene Ontology analyses did not reveal any enrichments in Cluster 1 (decreased in wild-type, increased in Δ*katG*) or Cluster 4 (decreased in wild-type, but not in Δ*katG*), several interesting processes emerged from Cluster 2 (increased in wild-type, decreased in Δ*katG*), Cluster 3 (increased in wild-type, but not in Δ*katG*), and Cluster 7 (increased in Δ*katG*, but not in wild-type). Cluster 2 was enriched for ATP synthesis and several stress response pathways. Cluster 3 was enriched for dicarboxylic acid biosynthesis, which includes the chorismate and folate biosynthesis pathways, and for tetrapyrrole biosynthesis, which includes much of the Vitamin B_12_ biosynthesis pathway. Cluster 7 was enriched for DNA damage and repair processes.

Because DNA damage and repair processes were up-regulated in BDQ-treated Δ*katG* cells but not BDQ-treated wild-type cells (Fig. 4f), we hypothesized BDQ induces increased DNA damage in Δ*katG* cells and Δ*katG* cells would therefore be hyper-susceptibility to further DNA damage. Consistent with this hypothesis were the increases in BDQ-induced 8-oxo-dG accumulation we had measured earlier (Fig. 3e). These have been shown to enable lethal double-strand DNA breaks following bactericidal antibiotic treatment in other bacteria^35^. To test this hypothesis, we treated wild-type and Δ*katG* H37Rv cells with the DNA damaging agent phleomycin and observed hyper-susceptibility in Δ*katG* cells (Fig. 4g). Similarly, mc^2^8245 cells were also hyper-susceptible to phleomycin over mc^2^7902 cells. These suggest the inability to tolerate BDQ-induced DNA damage may also help sensitize KatG-deficient cells to BDQ.

Collectively, these results indicate KatG deficiency confers BDQ hyper-susceptibility by altering several aspects of Mtb physiology, including synergistic repression of BDQ-induced *inhA* and *atpE* expression, epistatic induction of sensitizing transcriptional programs, and impaired DNA repair.

### Metabolic remodeling sensitizes KatG-deficient cells to bedaquiline

To further understand how KatG deficiency alters Mtb’s physiology, we performed genome-scale metabolic modeling analyses using our measured transcriptomic profiles as modeling constraints^36^. We utilized the most comprehensive model of Mtb H37Rv (iEK1011^37^) and generated a Δ*katG*-specific model by removing the catalase reaction. We then generated models corresponding to each of our 4 experimental sample conditions by applying the iMAT algorithm^38, 39^ to our RNA-sequencing data. We collected 10,000 flux samples for each metabolic reaction using the optGpSampler algorithm^40^ in the COBRA Toolbox^41, 42^, yielding metabolic flux distributions that quantitatively characterize the predicted activity for each metabolic reaction^36^ (Supplemental Table 4).

To assess model fidelity, we analyzed several metabolic reactions corresponding to our measured changes in ROS and to processes enriched in each of the transcriptomic clusters. Consistent with our observations that BDQ induces ROS accumulation (Fig. 3), model simulations predicted an increase in catalase activity in wild-type BDQ-treated over untreated cells (Fig. 5a), validating this modeling approach. Consistent with synergistic repression of *inhA* expression (Fig. 4c), model simulations predicted decreased mycolic acid biosynthesis (Fig. 5b). Model simulations also predicted decreased activity in propionyl-CoA metabolism (Fig. 5c), which has been shown to play important roles in mediating drug susceptibility in Mtb^30, 43^. Interestingly, although gene expression for *aroF* (Rv2540: chorsimate synthase), *folP1* (Rv3608c: dihydropteroate synthase) increased (Fig. 5d), model simulations predicted decreased metabolic activity through these enzymes and the rest of folate biosynthesis pathway (Fig. 5e). These results reveal epistatic suppression of folate biosynthesis in BDQ-treated *ΔkatG* cells that are not captured by the RNA sequencing data and suggest the induction of these genes might be a homeostatic response to deficiencies in their metabolic activities.

**Fig. 5.**
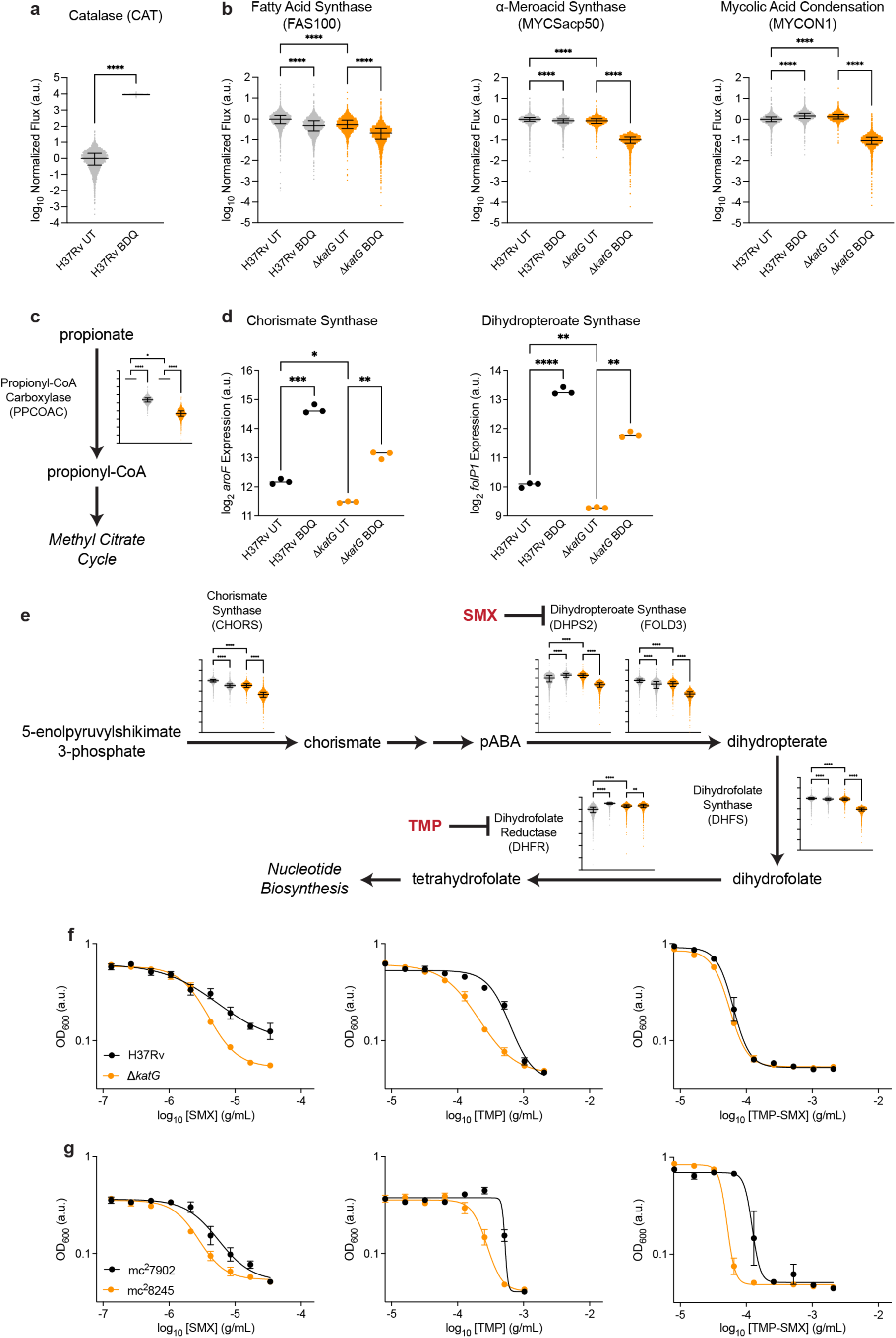
Metabolic remodeling sensitizes KatG-deficient cells to bedaquiline. **a**, Simulated catalase (iEK1011 CAT reaction) activity for BDQ-treated and untreated H37Rv cells from 10,000 samples using the iEK1011 Mtb genome-scale metabolic model. **b**, Simulated fatty acid synthase (FAS100), ɑ-meroacid synthase (MYCSacp50), mycolic acid condensation (MYCON1) activities for BDQ-treated and untreated H37Rv and Δ*katG* cells. **c**, Simulated propionyl-CoA carboxylase (PPCOAC) activity for BDQ-treated and untreated H37Rv and Δ*katG* cells. **d**, *aroF* and *folP1* expression is increased in BDQ-treated and untreated H37Rv and Δ*katG* cells. **e**, Simulated chorismate synthase (CHORS), dihydropteroate synthase (FOLD3 and DHPS2), dihydrofolate synthase (DHFS), and dihydrofolate reductase (DHFR) activities are synergistically decreased in BDQ-treated Δ*katG* cells. **f**, Δ*katG* cells are sensitized to SMX and TMP in 14-day dose-response experiments. **g**, mc^2^8245 cells are sensitized to SMX, TMP, and their combination in 14-day dose-response experiments. n = 3 biological replicates for each experiment. Brown-Forsythe and Welch ANOVA statistical tests were performed on RNA expression data with comparisons between BDQ-treated and untreated cells wild-type and Δ*katG* cells or between untreated wild-type and Δ*katG* as indicated with FDR correction. *: p ≤ 0.05, **: p ≤ 0.01, ***: p ≤ 0.001, ****: p ≤ 0.0001.

We previously demonstrated bactericidal antibiotics also induce nucleotide biosynthesis as a homeostatic response to stress-induced nucleotide pool disruptions^44^. Nucleotide metabolism is important for DNA replication and repair, which would be expected to increase given accumulation of oxidative DNA damage (Fig. 3e) and induction of DNA repair genes (Cluster 7; Fig. 4f). Consistent with our previous findings, model simulations predicted that BDQ treatment synergizes with KatG deficiency to suppress nucleotide metabolism, including downregulation of PRPP synthase and purine and pyrimidine biosynthesis activities (Extended Data Fig. 2).

Folate metabolism is essential for providing tetrahydrofolate substrates for *de novo* nucleotide biosynthesis. Because model simulations predicted decreased folate biosynthesis in Δ*katG* cells but not wild-type cells, we hypothesized Δ*katG* cells would also be hyper-susceptible to folate metabolism inhibition. To test this hypothesis, we performed dose response experiments in wild-type and Δ*katG* H37Rv cells and mc^2^7902 and mc^2^8245 cells using sulfamethoxazole (SMX), trimethoprim (TMP), and their combination. Dihydropteroate synthase is the target for SMX and dihydrofolate reductase is the target for TMP and these molecules are widely used as bacteriostatic antibiotics for several bacterial pathogens, but not Mtb. Consistent with our hypothesis, Δ*katG* and mc^2^8245 cells were hyper-susceptible to these folate metabolism inhibitors (Fig. 5f and 5g), indicating defects in BDQ-induced folate metabolism may also contribute to BDQ hyper-susceptibility in KatG deficient cells.

Collectively, these modeling suggest KatG deficiency confers BDQ hyper-susceptibility by also altering several aspects of Mtb metabolism, including epistatic inhibition of mycolic acid biosynthesis, propionate metabolism, and folte biosynthesis.

### Isoniazid resistant clinical strains are hyper-susceptible to bedaquiline

To evaluate the clinical relevance of these findings, we performed a meta-analysis of publicly available drug susceptibility data. Because *katG* loss-of-function mutations are the primary cause of INH resistance^8^, we hypothesized INH-resistant clinical isolates would possess lower BDQ minimum inhibitory concentrations (MICs) than INH-susceptible clinical isolates. The World Health Organization-sponsored Comprehensive Resistance Prediction for Tuberculosis: an International Consortium (CRyPTIC) consortium recently completed a large-scale curation and characterization of 12,289 Mtb clinical isolates from 23 countries in Asia, Africa, South America, and Europe and across 5 Mtb lineages^16^. Analyzing the CRyPTIC data, we found a global reduction in BDQ MICs at the population level between INH-resistant and INH-susceptible isolates (Fig. 6a). Interestingly, the CRyPTIC consortium predicted the probability of BDQ resistance in INH-resistant isolates would also be only 1.5%, lower than that of any of the other 11 drugs tested^16^.

**Fig. 6.**
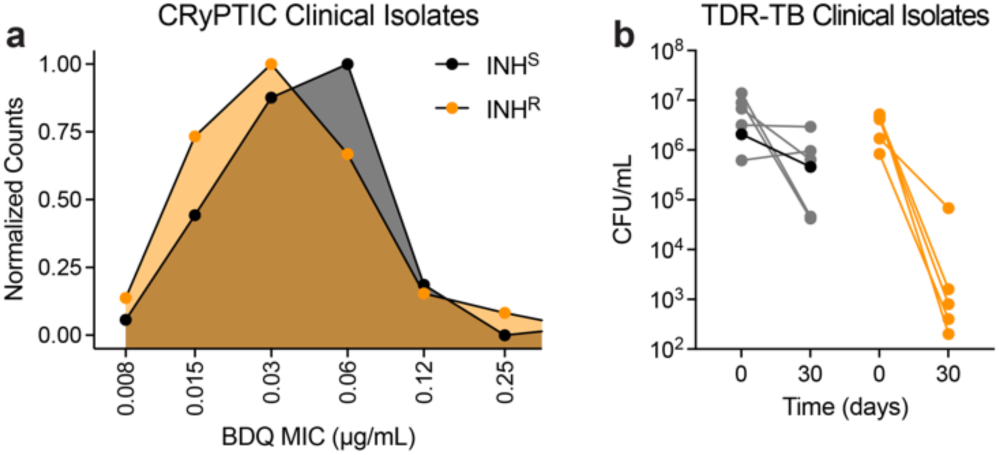
Isoniazid resistant clinical strains are hyper-susceptible to bedaquiline. **a**, Normalized count distributions of BDQ MICs in INH-susceptible and INH-resistant clinical strain data from the CRyPTIC Consortium^16^. **b**, INH-resistant clinical strains from the TDR-TB strain bank^17^ (orange: TDR-0019, TDR-0031, TDR-0042, TDR-0193, TDR-0198) have overall greater susceptibility to 2.7 μg/mL BDQ than INH-susceptible (gray: TDR-0077, TDR-0081, TDR-0126, TDR-0164) clinical strains in 30-day time-kill experiments. Wild-type H37Rv is included for reference (black). n = 1 biological replicate for each strain.

To validate these findings, we randomly selected 5 INH-resistant and 5 INH-susceptible clinical strains from the World Health Organization- and United Nations-sponsored Special Programme for Research and Training in Tropical Disease *M. tuberculosis* (TDR-TB) strain bank (Supplemental Table 5), which were curated in 2012 before the development and dissemination of BDQ as a TB therapeutic. We treated these cells will 2.7 μg/mL BDQ and enumerated CFUs after 30 days of BDQ treatment. We found these INH-resistant cells trended towards BDQ hyper-susceptibility over INH-susceptible cells (Fig. 6b). Importantly, 4 of the 5 INH-resistant cells we tested possess S315T mutations in KatG, the most clinically prevalent form of INH resistance^8^. Together these results indicate BDQ hyper-susceptibility in INH-resistant cells is a naturally occurring and clinically important phenotype which may explain the significant efficacy in using BDQ to treat MDR-TB.

## DISCUSSION

MDR-TB, defined as resistance to both first-line drugs INH and RIF, is a growing source of global mortality and morbidity and threatens global control of TB disease^1^. These have motivated several drug discovery efforts to identify efficacious anti-tubercular compounds for treating MDR-TB, resulting in the discovery and development of BDQ^4, 5^ which is now a cornerstone for several MDR-TB regimen including BPaLM^6^. Here we have shown the efficacy of BDQ against MTB-TB may be mechanistically explained by BDQ hyper-susceptibility in KatG loss-of-function mutations in INH-resistant cells, the dominant form of INH resistance. From our systems biology analyses, we propose KatG-deficiency induces BDQ hyper-susceptibility by several complementary physiological mechanisms, deficient capacity for detoxifying BDQ-induced ROS and repairing BDQ-induced DNA damage, synergistic transcriptional repression of INH and BDQ’s drug targets, induction of sensitizing transcriptional programs, and metabolic repression of mycolic acid, propionate, and folate biosynthesis. We further demonstrate BDQ hyper-susceptibility is common in INH-resistant clinical isolates curated by World Health Organization- and United Nations-sponsored programs, including INH-resistant strains containing KatG S315T mutations.

Here we report synergistic lethality between long-term INH and BDQ co-treatment. These results directly contradict observations by several groups that BDQ antagonizes INH efficacy^13–15^. However, each of these previous studies reached these interpretations by enumerating CFU after less than 2 weeks of co-treatment with INH and BDQ. Consistent with these reports, we also found antagonistic activity over a 2-week period. However, in contrast we found synergistic lethality in the 14-30 days after initiating INH and BDQ co-treatment. In both our study here and one of these previous studies^13^, INH mono-treatment CFUs cross over with BDQ co-treatment CFUs after 10-14 days of co-treatment. Our findings here reveal a biphasic response to BDQ co-treatment characterized first by BDQ suppression of INH lethality in the early phase, followed BDQ suppression of INH resistance in the late phase. These results are likely explained by times scale differences for INH and BDQ activity. INH’s activity is triphasic, involving an initial bacteriostatic period for the first 24 hours, followed by a rapid 3-4 log bactericidal period over 3-4 days, followed by rapid selection and development of INH resistance^8^. In contrast, BDQ’s activity is biphasic, involving an initial bacteriostatic period for the first 2-3 days, followed by a slow bactericidal period which does not achieve 3-4 log bactericidal activity until at least after 10 days of treatment^11^. Here we propose because INH requires NAD^+^ supplied by the respiratory chain as an oxidating agent to activate the INH prodrug^7^, BDQ’s inhibitory impact on the respiratory chain is likely to antagonize INH’s early bactericidal activity^13^, while BDQ’s slow activity induces late phase synergistic lethality.

Based on our observations that BDQ suppresses fixation of INH resistance, we propose INH resistance confers hyper-susceptibility to BDQ. To our knowledge, collateral sensitivity between INH resistance and BDQ has never been directly studied. A recent report involving laboratory evolution of the auxotrophic Mtb strain mc^2^6206 to 23 anti-tubercular compounds reported collateral sensitivity to BDQ in 2 of 3 INH-resistant strains selected^18^. Our work here supports these observations and provides mechanistic detail into this interaction. However, there exists some nuance to our findings here. First, although *katG* loss-of-function mutations are the most significant source of INH resistance, mutations in the promoter region of *inhA* and within other genes have also been shown to confer INH resistance^8^. In this present study, we focused on KatG deficiency and did not study these other forms of INH resistance, which we acknowledge may confer different phenotypes. Importantly, in the recent study involving mc^2^6206, all 3 INH-resistant strains possessed mutations in *katG*, but only 2 of 3 strains were hyper-susceptible to BDQ by MIC. Consistent with these, our analyses of CRyPTIC drug susceptibility data indicate although INH-resistant mutants possess decreased BDQ MICs at the population level, there exist several INH-resistant *katG* mutants that possess BDQ MICs similar to INH-susceptible strains. It is possible that experimental design and execution considerations may explain these discrepancies. However, it is also likely that in these laboratory-evolved and naturally-occurring clinical strains there also exist other genomic mutations that may compensate either directly for deficiency in KatG’s catalase activity (such as over-expression of *ahpC*^45^) or for other processes related to the mechanisms described here. These results underscore the need for more work into better understanding collateral sensitivity mechanisms in MDR-TB.

Several mechanisms have been proposed for how BDQ exerts its lethality, including ATP starvation^10, 11^ and respiratory uncoupling^12^. Although we did not directly test if KatG cells are hyper-susceptible to ATP depletion, we did not find sensitization to the respiratory uncouplers carbonyl cyanide m-chlorophenyl hydrazone (CCCP) and nigericin (Extended Data Fig. 3). It is therefore unlikely that BDQ-induced uncoupling explains BDQ hyper-susceptibility in INH-resistant cells. Because BDQ treatment induces oxidative stress regulons in mycobacteria^12^ and triggers increases in Mtb respiration^27^, BDQ has also been proposed to stimulate ROS production, which has been shown to participate in antimicrobial for Mtb and other bacteria^46–48^. However, BDQ-induced ROS formation has never been directly observed in Mtb and, in contrast, has been reported by others to not occur at all^27^. Here we report for the first time, to the best of our knowledge, that BDQ treatment is indeed sufficient for inducing ROS formation and that these are potentiated in KatG deficient cells. Several possible explanations may underlie discrepancies between our results here and previous studies. First, ROS are short-lived and highly sensitive to experimental conditions. The previous study that was unable to find BDQ-induced ROS^27^ had treated cells for 1 hour and incubated with ROS-sensitive dyes for 30 minutes before measuring dye fluorescence by flow cytometry. Here, we treated Mtb cells with BDQ for 4 days and incubated the cells with CellRox for 24 hours. It is possible 1-hour BDQ treatment or 30-minute CellRox incubation was insufficient for generating measurable changes. In addition, many ROS-sensitive dyes possess proprietary chemical structure which limits understanding into how these dyes are activated and with what specificity. There is consensus amongst the redox research community that multiple orthogonal assays should be performed to confidently assess ROS formation and oxidative cellular damage^49^. Here we assayed ROS and oxidative damage using 3 independent assays for ROS-sensitive dye oxidation, protein carbonylation, and guanine oxidation, each supporting the hypothesis that BDQ sufficiently induces ROS formation in Mtb. These results conclusively demonstrate BDQ treatment over a time period relevant to BDQ’s slow activity^11^ generates measurable ROS and highlights the importance of contextualizing data within experimental designs during interpretation.

Our findings here indicate many changes in Mtb physiology beyond deficient oxidative stress detoxification and DNA repair also contribute to BDQ hyper-susceptibility in KatG-deficient cells. We propose a model in which KatG deficiency induces several transcriptional changes that jointly sensitize INH-resistant cells to BDQ. These include genetic repression of drug target expression, induction of sensitizing transcriptional regulons, and epistatic repression of metabolic pathway enzymes. We speculate none of these physiological alterations alone are sufficient for fully sensitizing KatG-deficient cells to BDQ, but rather BDQ hyper-susceptibility is an emergent and epistatic property of several effects. For instance, it is possible down-regulated folate and nucleotide biosynthesis may synergize to limit DNA repair of BDQ-induced oxidative DNA damage.

Moreover, it is well-known that bacterial metabolism is important for antimicrobial efficacy^50^ and that INH is only effective against metabolically active Mtb cells^8^. While previous studies primarily focus on oxidative phosphorylation^13^, here we propose metabolic inhibition of mycolic acid and folate biosynthesis may also contribute to BDQ hyper-susceptibility in INH-resistant cells. Although we did not directly quantify mycolic acid biosynthesis rates, it is reasonable to hypothesize inhibition of mycolic acid biosynthesis by BDQ could phenocopy the effects of INH in INH-susceptible cells and elicit some lethality in KatG-deficient cells. In support of this hypothesis, previous meta-analyses reveal frequent co-occurrence of *katG* and *inhA* mutations in clinical Mtb strains^51^, suggesting possible selection for mycolic acid biosynthesis gain-of-function mutants to compensate for naturally occurring mycolic acid biosynthesis defects in KatG-deficient cells. We additionally propose BDQ-induced inhibition of several other biosynthetic pathways, including chorismate, folate, propionyl-CoA, and nucleotides might also contribute to BDQ hyper-susceptibility. Here we validated the potential contribution of folate biosynthesis defects to BDQ hyper-susceptibility using SMX, TMP, and their combination. Our findings suggest these inducing deficiencies may also mechanistically explain previous reports that TMP and SMX suppress emergence of INH resistance in Mtb^52^.

Our work here also highlights how systems biology approaches, including predictive genome-scale metabolic modeling^36, 53^, can support mycobacterial research and enable experimentally testable hypotheses^44^. It is important to note that metabolic modeling simulations predict decreased pathway activity despite increased gene expression for several important folate biosynthesis genes. These suggest these pathway activities are substrate-limited rather than enzyme-limited and demonstrate how gene expression changes alone may not always accurately predict how metabolic pathway activities are altered. Similarly, we observed significant lethality in our laboratory-evolved *katG^-^* cells (Fig. 2a) despite negligible changes in MIC (Fig. 2c). These highlight how important insights into drug-susceptibility and/or drug-resistance physiology may be hidden by traditional approaches in measuring MICs or performing RNA sequencing experiments, in support of recent observations that sub-breakpoint changes in MIC are sufficient for predicting TB reinfection after 6-months first-line therapy^54^. Indeed, efforts integrating transcriptomic^20, 21, 55^, metabolomic^56^, fluxomic^57^, lipidomic^58^, chemogenomic^22, 59–61^, and interpretable machine learning^44, 62^ approaches are needed to better understand how Mtb physiology constrains intracellular infection and drug efficacy^63–66^.

Finally, here we validated BDQ hyper-susceptibility in INH-resistant Mtb clinical isolates. These are important for establishing clinical relevance and for enabling therapeutic translation. Our findings here demonstrate how BDQ’s exceptional efficacy against MDR-TB^5^ can be mechanistically explained by BDQ hyper-susceptibility in KatG-deficient cells. Understanding such mechanisms are important for enabling new strategies to rationally designing MDR-TB treatment regimen^67, 68^, for instance by targeting Mtb *physiological* processes instead of only essential genes^59–61^. For example, while there is considerable enthusiasm in developing new drugs for targeting energy metabolism^66^, our results here suggest drug regimen including folate biosynthesis inhibition with SMX and/or TMP may also be efficacious against MDR-TB. These strategies are largely unexplored but provide exciting opportunities for global control of TB disease.

## METHODS

### Bacterial strains, growth conditions, reagents

Bacterial strains used in this study were wild-type *M. tuberculosis* H37Rv, a H37Rv KatG deletion mutant, a laboratory-evolved H37Rv KatG_mut_ E553K, a laboratory-evolved H37Rv MmpR5_mut_ G65R, recombinant H37Rv strains transformed with plasmids expressing the transcription factors genes *furA*, *kmtR*, *prpR* or *Rv3160c* under a tetracycline-inducible promoter^20–22^, the Mtb auxotrophic strains mc^2^7902 (H37Rv Δ*panCD ΔleuCD ΔargB*) and mc^2^8245 (Δ*panCD ΔleuCD ΔargB Δ2116169-2162530*)^28^, and Mtb strains from the TDR-TB strain bank^17^. All strains were cultured in Middlebrook 7H9 liquid media (Difco) supplemented with 10% oleic acid-albumin-dextrose-catalase (OADC; Difco) and 0.05% Tyloxapol (Millipore Sigma) at 37°C with shaking. Liquid cultures were sub-cultured on Middlebrook 7H10 solid media supplemented with 10% OADC and 0.2% glycerol (Acros Organics). Liquid and solid media for Mtb auxotrophic strains were additionally supplemented with 24 μg/mL L-pantothenate (Millipore Sigma), 50 μg/mL L-leucine (Millipore Sigma), 200 μg/mL L-arginine (Millipore Sigma), and 50 μg/mL L-methionine (Millipore Sigma). All experiments were performed in at least biological triplicate as indicated.

### Time-kill experiments

Frozen stocks of H37Rv or auxotrophic Mtb cells were inoculated 1:50 into 7H9 culture medium and grown to mid-log phase at an OD_600_ 0.5. Cultures were then back diluted 1:50 into fresh media containing drugs with 0, 0.17, 0.68, or 2.7 μg/mL BDQ (Adooq Bioscience) and/or 0.1 μg/mL INH (Millipore Sigma) as indicated. Cultures were incubated at 37°C with shaking, sampled at indicated time points, serially diluted in 7H9 media, and plated on 7H10 OADC agar plates. Plates were incubated at 37°C for 3-6 weeks after which colony forming units were enumerated.

### Whole-genome sequencing

Mtb colonies were picked from 7H10 agar plates and cultured in 7H9 medium to mid-log phase. Following incubation, cultures were transferred onto ice for 10 min, washed twice with ice cold PBS, and pelleted by centrifugation at 6000 × RCF for 5 min. DNA was extracted using the QIAamp DNA Mini purification kit (Qiagen) according to the manufacturer’s instructions. DNA concentrations were measured using a Thermo Fisher Nanodrop. DNA integrity was confirmed using an Agilent 2100 Bioanalyzer system. Sequencing libraries were prepared using the Illumina Nextera XT DNA Library Prep Kit as per manufacturer’s instructions. DNA sequencing was on an Illumina MiniSeq system with 30x coverage. Raw sequencing reads were aligned against the NCBI *M. tuberculosis* H37Rv reference genome (NC_000962.3) using *Bowtie 2*^69^. Once aligned, variants were called using *SAMtools*^70^.

### Drug susceptibility testing

Growth inhibitions experiments were performed on Mtb strains for each drug using the EUCLAST method^71^. Briefly, compound plates containing test drugs were prepared with 2-fold serial dilutions in 7H9 medium. Mtb cells were grown to mid-log phase, diluted 1:100 in 7H9, and 50 µL were dispensed into each well to achieve a final working volume of 100 µL per well. Drugs tested included BDQ, H_2_O_2_, phleomycin, SMX, TMP, CCCP, or nigericin as indicated. TMP-SMX was prepared at a 60:1 ratio as done previously^52^. Plates were incubated for 8-14 days at 37°C and OD_600_ absorbance was measured on a Biotek Synergy H1 or a BMG Labtech PHERAstar microplate reader at 7, 8, or 14 days as indicated. For experiments involving inducible *furA*, *kmtR*, *prpR* or *Rv3160c* expression, cells were cultured in the presence or absence of 100 ng/mL anhydrotetracycline.

### Reactive Oxygen Species (ROS) quantification

ROS accumulation was measured using the ROS-sensitive dye CellRox Green (Invitrogen) according to the manufacturer’s instructions. Briefly, Mtb cells were grown to mid-log and back-diluted to OD_600_ 0.2. 3 mL cultures were incubated for 4 days with and without 0.68 µg/mL BDQ. Cells were dispensed into 96-well plates and incubated with 5 µM CellRox Green for 24 hr at a final working volume of 200 µL per well. CellROX fluorescence was measured on a Biotek Synergy H1 microplate reader at 520 nm.

### Protein carbonylation quantification

Protein carbonylation was measured using the OxiSelect Protein Carbonyl ELISA Kit (Cell Biolabs) according to the manufacturer’s instructions. Briefly, Mtb cells were grown to mid-log and cultured with and without 5.4 µg/mL BDQ at a final working volume of 10 mL for 16 hr. Following incubation, cultures were transferred onto ice for 10 min, washed twice with ice cold PBS, and pelleted by centrifugation at 6000x RCF for 5 min. 200 µL Bacterial Protein Extraction Reagent (B-PER II) was added to each pellet with 100 µg/mL lysozyme and 5 U/mL DNase I added. Total protein abundance was quantified for each sample using a Bicinchoninic acid (BCA) assay (Thermo Fisher Scientific) according to the manufacturer’s instructions before performing protein carbonylation experiments with the OxiSelect ELISA assay. Absorbance measurements were taken at 450 nm on a Biotek Synergy H1 microplate reader.

### 8-oxo-dG quantification

Protein carbonylation was measured using the OxiSelect 8-OHdG Oxidative DNA Damage ELISA Kit (Cell Biolabs) according to the manufacturer’s instructions. Briefly, Mtb cells were grown to mid-log and cultured with and without 2.7 µg/mL BDQ at a final working volume of 10 mL for 16 hr. Following incubation, cultures were transferred onto ice for 10 min, washed twice with ice cold PBS, and pelleted by centrifugation at 6000 × RCF for 5 min. DNA was extracted using the QIAamp DNA Mini purification kit (Qiagen) according to the manufacturer’s instructions. Double strand DNA was converted to single strand DNA by incubating samples at 95°C for 5 min and rapidly chilling on ice. Single strand DNA was then digested before performing 8-OHdG measurements using the OxiSelect ELISA kit. Absorbance measurements were taken at 450 nm on a Biotek Synergy H1 microplate reader.

### RNA Sequencing and Analyses

Mtb cells were grown to mid-log phase and cultured with and without 2.7 µg/mL BDQ for 16 hr. Immediately following incubation, cultures were pelleted and 1 mL Trizol (Thermo Fisher Scientific) was added to each pellet. Samples were lysed by bead beating and kept at 4°C for 5 min. Chloroform (Amresco) was added to each sample, lysates were vortexed vigorously, and samples were pelleted by centrifugation at 12000 × *g* for 5 min. Supernatants for each sample were carefully transferred to RNase-free tubes for purification by the Direct-Zol RNA Miniprep Plus RNA purification kit (Zymo Research), according to the manufacturer’s instructions. Total RNA was eluted in RNase free water and RNA concentrations were measured using a Thermo Fisher Nanodrop One. RNA integrity was confirmed using an Agilent 4200 TapeStation system. Ribosomal RNA was depleted using the Illumina Ribo-Zero Plus rRNA Depletion kit followed by library preparation with the NEBNext Ultra II Directional RNA Library Prep Kit for Illumina (New England Biolabs) as per manufacturer’s instructions. RNA sequencing was performed by the Rutgers Genomics Center on an Illumina NovaSeq6000 system with 100x coverage.

Raw sequencing reads were aligned against the NCBI *M. tuberculosis* H37Rv reference genome (NC_000962.3) using *Bowtie 2*^69^. Read counts were compiled using *featureCounts*/^72^ and quantile normalized by *qsmooth*^73^. Quality data, adapter and quality trimming statistics, and alignment and counts metrics were compiled and assessed using *MultiQC*^74^. Normalized gene counts were hierarchically clustered for clustering analyses.

### Gene set enrichment analyses

Gene set enrichment analyses were performed on Biocyc^75^ using the *M. tuberculosis* H37Rv curated database (v. 27.1), as previously described^44^. SmartTables were created for genes comprising each cluster set as determined by hierarchical clustering. Gene Ontology enrichment terms were identified by performing “Enrichment Analysis” for “GO terms – genes enriched for GO (biological processes) using Fisher Exact statistics and Benjamini-Hochberg corrections for false discovery.

### Genome-scale metabolic modelling

Model simulations were performed using the iEK1011 genome-scale model of *M. tuberculosis* metabolism^37^, as previously described^44^. RNA sequencing data were applied as modeling constraints using the *iMAT* algorithm^38, 39^ for sampling in the *COBRApy* toolbox^41^ with Gurobi Optimizer as the solver. KatG-deficient cells were modeled by setting bounds for the catalase (CAT) reaction to 0 to represent *katG* deletion. Simulations were performed by sampling each model 10,000 using *optGpSampler*^40^. Samples were down-sampled 10-fold for visualization purposes.

### WHO CRyPTIC MIC data analyses

BDQ minimum inhibitory concentration (MIC) data was downloaded from the WHO CRyPTIC Consortium^16^. Samples with missing BDQ MICs were removed from downstream analysis. BDQ MICs outside the reported concentration rate were replaced with adjacent concentrations on a dilution series as appropriate (MICs “≤ 0.008” set to “0.008”; “≤ 0.015” set to “0.015”; “> 1” set to “2”; “> 2” set to “4”). Samples were then segregated to INH-susceptible (INH^S^) or INH-resistant (INH^R^) as determined by the CRyPTIC Consortium. Strains were enumerated for each set and MIC and normalized by peak counts for each distribution (INH^S^ counts normalized to counts at 0.06 μg/mL BDQ; INH^S^ counts normalized to counts at 0.03 μg/mL BDQ).

## DATA AVAILABILITY

All data are available in the main text or supplementary materials. RNA sequencing data is available in on the Gene Expression Omnibus (GEOXXXXXX) and Sequence Read Archive (SRAXXXXXX).

## CODE AVAILABILITY

Analysis code is deposited on GitHub (https://github.com/jasonhyang).

## ACKNOWLEDGEMENTS

*Mycobacterium tuberculosis* H37Rv was generously shared by Deborah Hung from the Broad Institute of MIT and Harvard. H37Rv Δ*katG*, mc^2^7902, and mc^2^8245 were generously shared by William Jacobs from the Albert Einstein College of Medicine. Transcription factor over-expression strains were generously shared by David Sherman at the University of Washington. TDR-TB strains were generously shared David Alland at Rutgers New Jersey Medical School. The authors thank David Alland, Hassan Safi, and Pradeep Kumar, Joel Freundlich, and Jees Sebastian from Rutgers New Jersey Medical School; and David Sherman from the University of Washington for helpful discussions and experimental suggestions. The authors also thank James Gomez and Zohar Bloom-Ackermann from the Broad Institute of MIT and Harvard for training in Biosafety Level-3 research activities. This work was supported by grants R00-GM118907, U19-AI11276, U19-AI62598, and R01-AI146194 from the National Institutes of Health; HDTRA12210032 from the Defense Threat Reduction Agency; Rutgers New Jersey Medical School; the Broad Institute of MIT and Harvard; and a generous gift from Anita and Josh Bekenstein. J.H.Y. is additionally supported by the Agilent Early Career Professor Award.

## AUTHOR CONTRIBUTIONS

J.H.Y and J.J.C. conceptualized the study. J.H.Y., J.J.C., and S.M. supervised the project. N.O., M.H., M.L.C., A.S., B.L., S.L., S.M., and J.H.Y. executed the experiments and data analyses. N.O. and J.H.Y. wrote the manuscript and visualized the data. J.H.Y. and J.J.C. acquired funding. All authors reviewed the draft and assisted in manuscript preparation.

## EXTENDED DATA FIGURES

**Extended Data Fig. 1.**
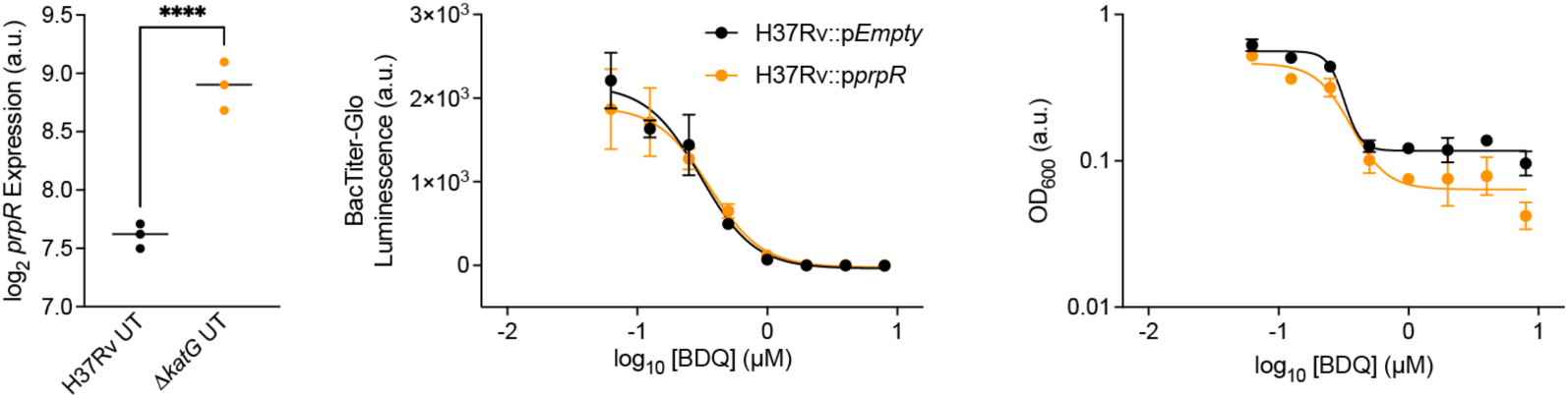
PrpR over-expression does not sensitize Mtb to BDQ. RNA sequencing reveals BDQ-induced up-regulation of *prpR* expression. *prpR* over-expression does not sensitize Mtb to BDQ over 7-day treatment as measured by either BacTiter-Glo or optical density. Smooth quantile normalized expression data for *prpR* given on a log_2_ scale. n = 3 biological replicates for each gene. Welch’s *t-*test was performed for comparisons between BDQ-treated and untreated cells wild-type cells. ****: p ≤ 0.0001.

**Extended Data Fig. 2.**
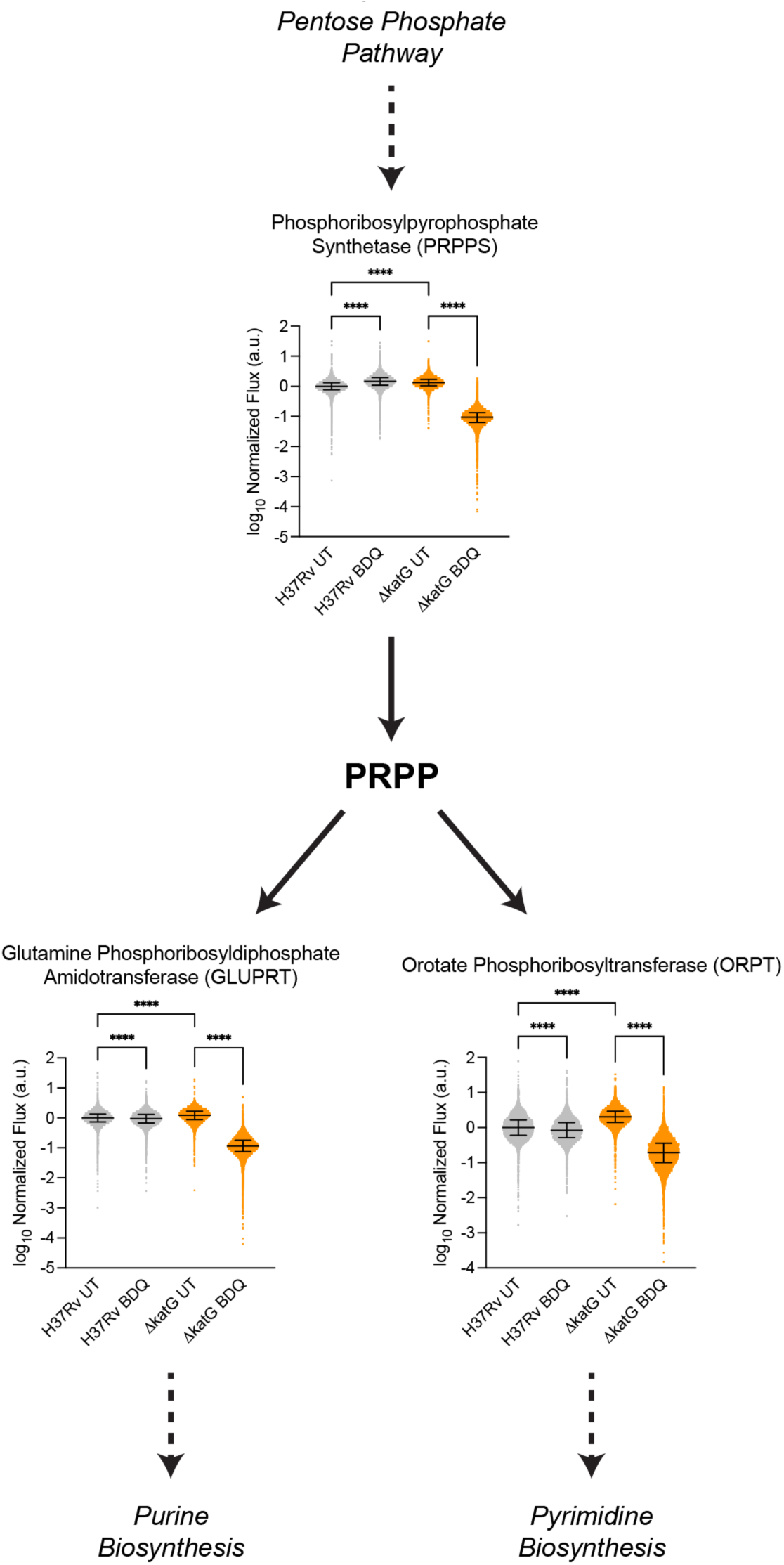
Bedaquiline synergistically inhibits nucleotide biosynthesis in Δ*katG* cells. Simulated phosphoribosylpyrophosphate synthase (iEK1011 PRPPS reaction), glutamine phosphoribosyldiphosphate amidotransferase (GLUPRT), and orotate phosphoribosyltransferase (ORPT) activities for BDQ-treated and untreated H37Rv cells from 10,000 samples using the iEK1011 Mtb genome-scale metabolic model. Brown-Forsythe and Welch ANOVA statistical tests were performed with comparisons untreated wild-type and Δ*katG* cells, BDQ-treated and untreated cells wild-type cells, and BDQ-treated and untreated Δ*katG* cells. ****: p ≤ 0.0001.

**Extended Data Fig. 3.**
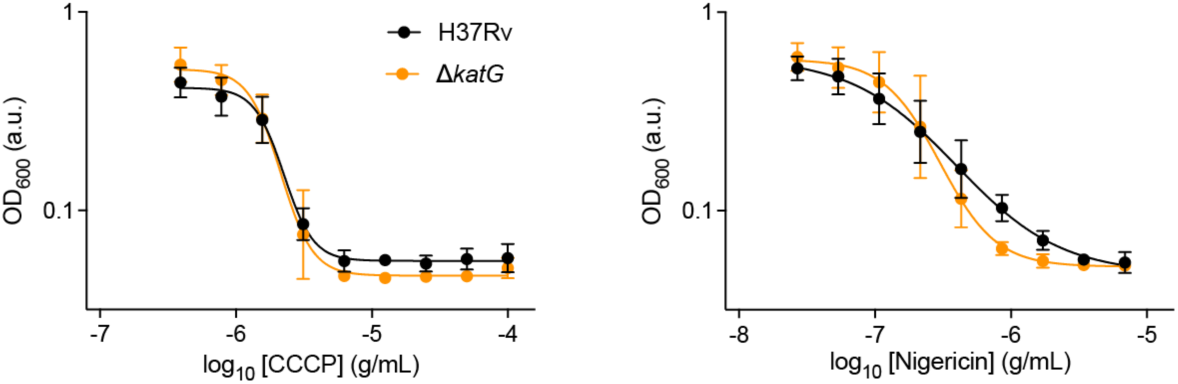
Δ*katG* cells are not hyper-susceptible to ionophoric stress. Wild-type and Δ*katG* H37Rv cells were treated with the ionophores carbonyl cyanide m-chlorophenyl hydrazone (CCCP) and nigericin in dose-response experiments for 14 days. n = 3 biological replicates for each experiment.

## SUPPLEMENTAL TABLES

**Supplemental Table 1. Mutations found in laboratory-evolved INH-resistant and BDQ-resistant *M. tuberculosis* H37Rv strains.**

**Supplemental Table 2. Smooth quantile normalized RNA-sequencing gene expression profiles.**

**Supplemental Table 3. Hierarchically clustered genes from BDQ-treated and untreated Δ*katG* and wild-type H37Rv cells.**

**Supplemental Table 4. iEK1011 Metabolic modeling simulations.**

**Supplemental Table 5. Strain information for isoniazid resistant and susceptible TDR-TB strains used in this study.**

**Supplemental Table 6. List of strains used in this study.**

